# The synaptic tag as an emergent biophysical state depending on the synaptic actin dynamics and the postsynaptic density

**DOI:** 10.1101/2025.02.18.638822

**Authors:** Francesco Negri, Jannik Luboeinski, Christian Tetzlaff, Michael Fauth

## Abstract

According to the synaptic tagging-and-capture hypothesis, long-term synaptic plasticity requires synapses to establish a synaptic tag, enabling them to capture plasticity-related products synthesized elsewhere in the neuron. The biophysical processes, molecules or properties that implement the tag are, however, unclear. We hypothesize that a mismatch between dendritic spine volume and the size of the postsynaptic density (PSD) inheres the essential characteristics of a synaptic tag. To test this, we developed a computational model that integrates established calcium-dependent synaptic plasticity with the complex biochemical dynamics of the PSD and actin – a key structural protein determining dendritic spine geometry. This model can explain a broad range of experimental findings across diverse synaptic plasticity protocols at the level of individual synapses as well as heterosynaptic plasticity protocols involving two synapses. Furthermore, the model predicts that repeated induction of plasticity within one hour results in a nonlinear accumulation of synaptic changes, reminiscent of the spacing effect observed in psychological studies of learning and memory. Hence, our model offers a concise mechanistic framework for understanding critical synaptic processes and demonstrates the viability of the hypothesis that temporal disparities in structural and biochemical dynamics form a memory trace that could act as a synaptic tag.

## Introduction

Synapses and their adaptation through synaptic plasticity are associated to the mechanisms of learning and memory [1]. The most prominent forms of synaptic plasticity are long-term potentiation (LTP, [2]) and long-term depression (LTD, [3]). Both have been found to undergo at least two major phases [4]: an early phase, decaying after a few hours, and a late phase that persists for several hours or even days. According to the *synaptic tagging-and-capture hypothesis*, the late phase only occurs if two factors are present at the same time at the (post)synapse [5, 6]: (i) plasticity-related products (PRPs) such as *de novo* synthesized proteins that can be captured by the synapse, and (ii) a so-called synaptic tag that indicates whether the synapse has received plasticity-triggering stimulation. The presence of the synaptic tag at the synapse then leads to the capture of PRPs. Note that both factors are present independently of each other, as a synaptic tag can be set without explicitly triggering PRP synthesis [5] or a tagged synapse can express late-phase plasticity by capturing PRPs that are being synthesized in response the stimulation of other synapses [7–10]. While numerous studies have investigated the trafficking, synthesis and turnover of plasticity-related proteins [11–14], the synaptic tag still remains a mystery. Despite experimental evidence suggesting that the synaptic tag is unlikely to consist of a single type of molecule [15] and instead reflects a transient structural state of the synapse [6], the specific biochemical or biophysical processes that form the synaptic tag are still unknown.

Most computational models characterizing the processes of the synaptic-taggingand-capture hypothesis are phenomenological [16–21], usually describing the synaptic tag as abstract variable that depends on the early-phase of synaptic plasticity. In this study, drawing from previous theoretical models [20, 22, 23], we develop a new tagging-and-capture model that captures the dynamic changes of the volume of the postsynaptic spine, which is based on the different dynamic states of the structural protein actin as well as of the postsynaptic density (PSD). Using this model, we test the hypothesis that a mismatch between actin-dependent spine volume and the PSD could represent the synaptic tag.

Actin is a globular protein that forms filaments. These preferentially polymerize at one end and depolymerize at the other, resulting in a continuous treadmilling. When close to the membrane, this induces pushing forces and retrograde movement of the filament [24], which continuously alters the spine shape [22, 23, 25, 26]. Additionally, multiple actin-binding proteins, such as the branching protein complex ARP2/3 or the severing protein cofilin, organize the filaments into continuously growing and reorganizing, branched networks. On the other hand, crosslinking proteins such as drebrin, actinin or CaMKII*β* [27, 28] can bind to the filaments, slowing down treadmilling and reorganization. This gives rise to at least two main populations of actin, exhibiting different treadmilling speeds [29], which we refer to as the *dynamic* and the *stable actin pool*. Typically, the activity of the dynamic actin pool is localized to discrete polymerization foci at the tip of the spine [30], where an increased retrograde movement of filaments is observed, whereas the stable actin pool forms a crosslinked mass at the bottom of the spine [29]. Following a plasticity-inducing stimulation, the concentrations as well as the reaction rates of actin-binding proteins in the spine undergo changes [28, 31]. While these changes are not long-lasting, they give rise to a rapid remodeling of spine geometry and alterations in the stable and dynamic actin pools [22, 26, 32, 33], which may persist on timescales of hours [22, 34].

To relate the dynamics of actin and the spine volume to long-lasting plasticity of the synaptic transmission efficacy, we focus on the dynamical behavior of the PSD. The PSD size (as measured by fluorescence or electron microscopy) is strongly correlated with the synaptic transmission efficacy (e.g., as measured by EPSPs [35]) through the number of AMPA receptors anchored at the postsynaptic site. In particular, it has been postulated that depending on its size, the PSD provides a number of slots for AMPA receptors [6, 36]. Moreover, in the basal state, the PSD size is correlated with the spine volume [37, 38], and previous modeling has shown that the PSD size determines the the stationary volume around which a spine fluctuates [23]. Thus, for each PSD size, there is a corresponding spine volume expected under basal conditions. Upon LTP, spines deviate from this basal correlation line and move towards higher volumes [38]. But then, they increase their PSD size and again decrease their volume, thereby reestablishing the basal correlation. Accordingly, on the population level, it has been observed that shortly after LTP, spines tend towards higher volumes and then relax back to the basal state after around 2 hours [31, 39, 40]. Taken together, there is a transient mismatch between spine volume and PSD, which is removed by adapting both volume and the PSD. In line with our hypothesis, this supports the view that the PSD is able to react to changes in actin-dependent spine volume and, when free PRPs are available in the spine, adapt its size to match the new spine volume.

In this study, we test the viability of our hypothesis using a mathematical model of actin-dependent volume dynamics and its interplay with PSD changes. After providing a detailed description of the core principles of the actin-PSD model, we first show in a proof-of-concept setup that the temporal development of the different biochemical and -physical processes can account for the formation of a synaptic tag on the experimentally observed timescale. Next, integrating the actin-PSD model with a more detailed model of calcium-dependent plasticity, we demonstrate the viabilityof our hypothesis in a situation comparable to the electrophysiological experiments. Further, we show for both versions of the model that the hypothesis also applies to multi-synaptic stimulation protocols, and that it can further provide new predictions about the synaptic dynamics given sequential stimulation protocols.

## Results

### Model for late-phase plasticity

Our proof-of-concept actin-PSD model (see Fig. 1) is based on recent insights that the spine volume results from the overall amount of actin filaments in a dynamic and a stable pool [22]. Here, we simplify the complex interplay between these two actin pools and the spine geometry by directly considering that the total spine volume *V*_tot_ is the sum of volumes resulting from the dynamic actin pool *V*_d_ and the stable actin pool *V*_s_. On the other hand, experiments show that there is a correlation between the spine volume and the size of the corresponding PSD [37, 38], and biophysical models demonstrate that a certain PSD size entails a certain spine volume even in the absence of actin [23, 26]. Therefore, we consider here a volume *V*_PSD_ to model the PSD size, which allows us to mathematically relate and compare it to the different actin volumes. In the following, we will describe the core biochemical and -physical processes influencing the dynamics of these different spine-related volumes.

**Fig. 1.**
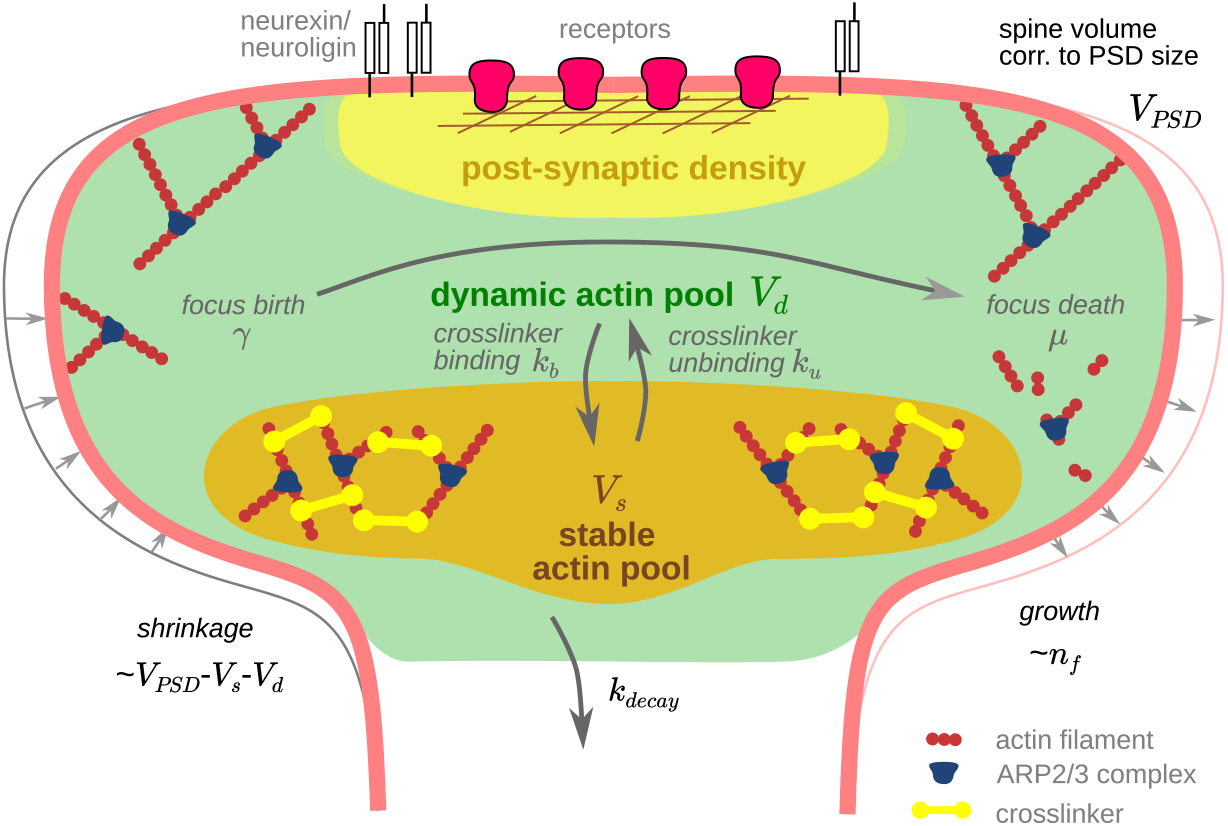
Sketch of the main quantities and processes in the actin-PSD model. The formation of foci leads to the generation of new actin filaments. These actin filaments are contained either in the dynamic or in the stable pool and can transition between these pools by binding and unbinding of crosslinkers. The total spine volume, which is a sum of the volumes from dynamic pool *V*_d_ and stable pool *V*_s_, is assumed to increase depending on the number of active actin polymerization foci *n*_f_ (right side) and shrinks, when it is larger than the volume expected for the current PSD size *V*_PSD_ (left side).

The temporal evolution of the volume resulting from the dynamic actin pool is determined by (i) the continuous transition of actin between dynamic and stable state due to crosslinker binding and unbinding, (ii) the formation of new actin filaments dependent on the number of polymerization foci, and (iii) counteracting forces from the cell membrane (cf. Eq. 1 in Methods):

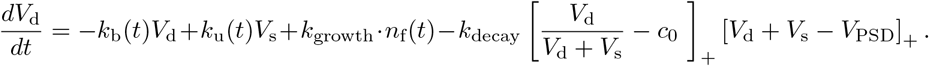

The first two terms on the right-hand side cover exchange between the stable and the dynamic pool determined by crosslinker binding with rate *k*_b_(*t*) and unbinding with rate *k*_u_(*t*). These rates are usually constant, but may change upon stimulation [27, 28, 41]. In general, the polymerization of new actin filaments happens in distinct polymerization foci [30], which are formed and dissolved stochastically [23]. We therefore describe the number of foci *n*_*f*_, which determines the third term on the right-hand side, by a birth-death process with birth rate *γ*(*t*) and death rate *µ*(*t*). Finally, the fourth term describes the mismatch between actin and PSD-dependent volumes: If the total synaptic volume *V*_tot_ = *V*_d_ + *V*_s_ is larger than the volume corresponding to the PSD size *V*_PSD_, the membrane exerts counteracting forces. These forces hinder actin polymerization or push material out of the spine, leading to a net reduction in the dynamic actin pool and the related volume *V*_d_ with rate *k*_decay_. Thus, the dynamic pool shrinkage is considered to be proportional to the difference between actin and PSD volume (as long as it is positive). If there is a larger dynamic actin volume, this effect will be stronger. Hence, the above described volume-dependent shrinkage is scaled by a factor that describes by how much the fraction of dynamic actin exceeds an offset concentration *c*_0_.

The evolution of the spine volume that results from the stable pool is fully described by the continuous exchange of actin between the stable and the dynamic pool (cf. Eq. 2 in Methods):

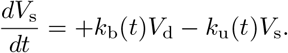

The dynamics of the PSD, and hence of *V*_*P SD*_, is gated by the availability or concentration of PRPs, represented by *ϕ*(*t*), and it will only change if the current spine volume *V*_tot_, resulting from the actin-dynamics, is not in equilibrium with the PSD-dependent volume. The temporal evolution is therefore given by (cf. Eq. 3 in Methods):

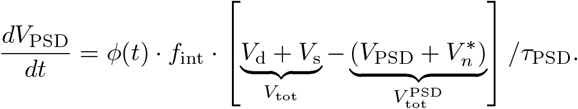

Here, 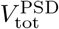 is the total spine volume expected for the current PSD size given the stochastic polymerization of actin filaments (*n*_f_(*t*)), which leads to the correction factor 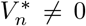 (see Methods). The factor *f*_int_ accounts for the physical scaling of the integration of PRPs into the PSD. It can be seen that when PRP is available (*ϕ* = 1), the volume corresponding to the PSD size will evolve to reestablish the equilibrium with the actin-induced volume. Thus, given our hypothesis that the temporal difference between actin-dependent spine volume and PSD size could represent the synaptic tag, we define in the following the synaptic tag as 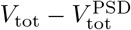.

In the following, we show (1) how this tag can be set by modulation of actin dynamics, (2) how such modulations are induced by neural activity, (3) that the tag can account for heterosynaptic plasticity and that it decays on a similar timescale as observed in experiments, and (4) that our tag model can explain and predict the temporal interaction of multiple stimuli at the same synapse.

### Setting the tag by modulation of actin dynamics

As a first step, we show that stimulus-induced changes in actin dynamics in our actin-PSD model can lead to the formation of a synaptic tag, which results in synaptic dynamics comparable to experimental findings. To this end, we test how our late-phase plasticity model behaves under standard plasticity-inducing stimuli, i.e., tetanus stimuli that induce LTP, and low-frequency stimuli (LFS) that induce LTD (cf. [44]). We model the influence of these stimuli directly by changing the parameters governing the actin dynamics. Thus, we switch from the basal actin parameter values to plasticity-related values during stimulation and back to the basal values after stimulation (see Methods and bottom of panels in Figs. 2-5 for the respective time intervals). In particular, the following parameters of the actin dynamics are changed (see Table 2): First, it is known that during LTP, crosslinkers rapidly unbind from the actin filaments, filaments are severed, and new actin foci emerge [28]. Similarly, also during LTD, crosslinker dynamics are altered [41, 45, 46]. In our model, this is reflected by a change in the crosslinker binding and unbinding rates *k*_b_ and *k*_u_. Second, the influx of calcium during both LTD and LTP triggers multiple signaling chains, which regulate, inter alia, the actin severing protein cofilin [47]. This can lead to increased severing and thus death of actin polymerization foci, but also to a large concentration of free actin monomers and nucleation of new filaments. Thus, in the model, we vary birth and death rates of actin foci accordingly, resulting in changes of *n*_f_.

**Fig. 2.**
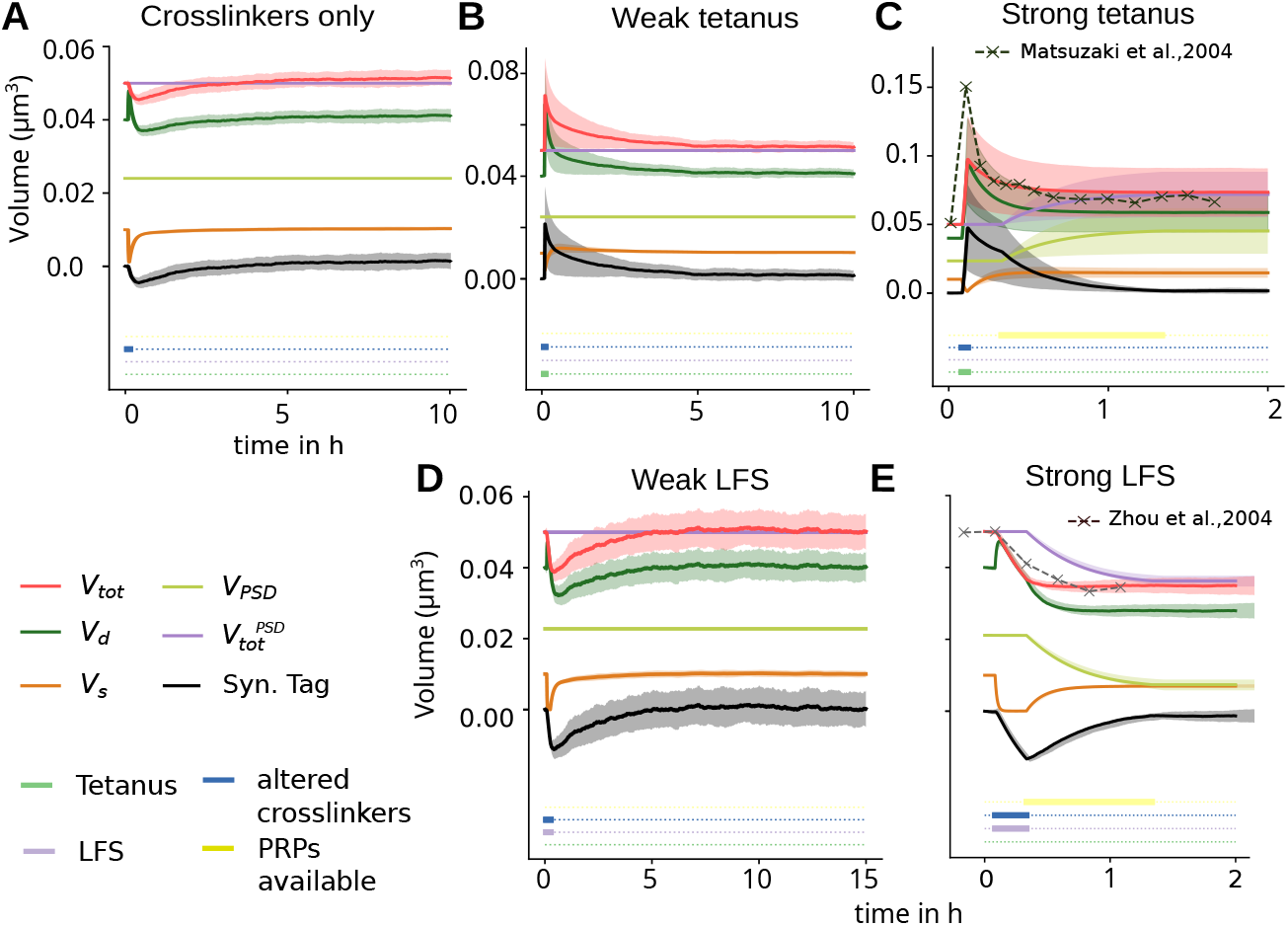
The dynamics of the proof-of-concept model given standard LTP and LTD protocols. Time evolution of the hypothesized synaptic tag (black), the dynamic and stable actin volume, the PSD volume *V*_PSD_, and derived quantities (*V*_tot_ = *V*_d_ + *V*_s_ and 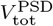) over time under different conditions: (A) Plasticity-related alteration of crosslinker (un)binding (interval indicated by dark blue bar at bottom); (B) a weak tetanus (green bar) without PRP synthesis; (C) strong tetanus with PRP synthesis (yellow bar); (D) low-frequency stimulation (purple bar) without PRP synthesis and (E) with PRP synthesis. Lines and shaded areas represent mean and standard deviation over 100 simulations, respectively. See Methods for details on the used stimuli. Dashed lines in panels C and E mark experimental results from [42] and [43].

To assess the influence of individual parameter changes, we first tested what would happen if only the crosslinker binding and unbinding rates (*k*_b_, *k*_u_) are transiently altered (Fig. 2A). As expected, we found that this leads to a rapid transfer of actin from stable (ocher curve) to dynamic pool (dark green curve) and the subsequent slow restoration of the stable pool to its stationary value, whereas the total volume of the actin pools only is transiently lowered and then quickly returns approximately to the basal level (pink curve). As the PSD volume does not change, the proposed synaptic tag (black curve) mirrors this behavior. Hence, altered crosslinkers alone will not induce long-term plasticity.

On the other hand, when the crosslinker dynamics (*k*_b_, *k*_u_) are altered and the birth rate of new foci (*γ*) is increased in parallel (Fig. 2B), which corresponds to a weak tetanization in experiments, an increase in the total volume is elicited. As compared to the case with crosslinkers only, we first observe an even stronger increase of the dynamic pool, which induces a slight and transient increase of the stable pool above its stationary level after the crosslinkers return to their normal rates. However, without PRP synthesis all quantities ultimately return to their initial (stationary) values and the proposed tag (black) decays. This is because the decay term in *V*_d_ drives the total volume of actin towards the volume determined by the PSD size *V*_PSD_. However, once an increase in PRP synthesis is present (as during strong tetanization), *V*_PSD_ and, thus, 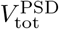 become dynamic and adapt to the perturbed total actin volume (Fig. 2C, violet and ocher). This leads to an accelerated decay of the proposed synaptic tag. As soon as 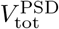 reaches the total actin volume, no further change happens, and the volume modifications become permanent. Note that in the presence of active PRP synthesis the resulting stable state 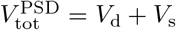 is a line attractor, which enables the total volume to take values from a continuous range (cf. Appendix). Further note that the parameter values were chosen such that the time course of *V*_tot_ resembles the relative volume changes from LTP-inducing glutamate uncaging experiments ([42], black dashed line in Fig. 2C).

Similar to the workings of LTP, our model can also account for LTD. Following experimental data, LTD yields the unbinding of crosslinkers, but in contrast to LTP the birth rate of new foci is decreased. These two changes together lead to a smaller increase in the dynamic pool volume upon stimulation and a decrease in the total actin volume, as well as a negative synaptic tag (Fig. 2D-E). When no PRP synthesis occurs, the total volume starts to increase again as soon as the stimulus ends and the birth rate of new foci is restored until the pre-stimulus level is reached (Fig. 2D). Once PRP synthesis is increased, the negative value of the tag causes a gradual decrease in *V*_PSD_, which in turn removes the tag (Fig. 2E). Note that parameter values have been chosen such that the model matches the time course of spine volumes after low-frequency stimulation observed in experiments (dashed line, [48]).

In summary, the proposed actin-PSD model can express both early and late LTP as well as early and late LTD and reproduces the respective experiments using one and the same parameter set. This indicates that the difference between actin-dependent spine volume and PSD size sets a synaptic tag such that synaptic changes can be consolidated into long-lasting plasticity.

### Integration with calcium-based early-phase plasticity and synthesis of plasticity-related products (PRPs)

As a next step, we investigate how the above described modulations in actin dynamics directly interact with pre- and postsynaptic spiking activity, calcium-dependent plasticity, and PRP synthesis dynamics. For this, we integrate our actin-PSD model with a calcium-based early-phase plasticity model (cf. [49]) and a PRP synthesis model (cf. [16]), which are well established from earlier studies on synaptic tagging and capture (see [16, 18–20]). We refer to this in the following as the “full-integrated model”. This model comprises a weight component for early-phase plasticity that undergoes LTD when the calcium amount exceeds a threshold value *θ*_d_, and LTP when calcium exceeds an even higher threshold *θ*_p_. Otherwise the early weight decays towards its baseline value zero. PRP synthesis takes place when the deviation from the baseline, summed over all synapses of a neuron (referred to as “signal triggering PRP synthesis”, SPS, cf. Eq. 6) exceeds a threshold *θ*_*ϕ*_.

In the full-integrated model, our actin-PSD late-phase plasticity model is also coupled to neuronal activity, assuming that the calcium level at the postsynaptic site determines the crosslinker activity and the dynamics of actin foci. When calcium is above the LTD threshold *θ*_d_, the signaling chain for depression is activated. We here assume calcineurin (CaN, [43, 47]) as a representative of this signaling chain and include a calcineurin signal that activates when calcium is above that threshold. The signal remains activated for a certain time afterward and as long as it is active, the above described effects of the LFS on the late-phase model are applied. Similarly, for tetanic stimulation and LTP, we model a CaMKII signal that triggers the proposed effects of tetanic stimulation on the late-phase model. This signal is activated when calcium exceeds an even higher threshold *θ*_CaMKII_.

Given the direct relation between the PSD size, corresponding to *V*_PSD_, and the number of AMPARs, we assume that the late-phase weight *z* is a linear function of *V*_PSD_ (cf. Eq. 8). This late-phase weight is added to the early-phase weight *h* to yield the total weight of the synapse as *w*(*t*) = *h*(*t*) + *z*(*t*).

Using this model together with leaky integrate-and-fire neurons, we could induce plasticity via a presynaptic spiking activity resembling the stimulation used in classical tagging-and-capture experiments (cf. [44]; see Methods), similar as for the simple switching of the rate parameters in our late-phase-only model earlier.

During a weak tetanus (Fig. 3A), calcium rises above the thresholds for early LTP and CaMKII activation. This leads to similar actin dynamics as in Figure 2B. The induced plastic changes are, however, not large enough for the SPS signal to cross the threshold for PRP synthesis (Fig. 3A bottom), such that all quantities decay to their initial values. The strong tetanus (Fig. 3B), which consists of three prolonged stimulation intervals of the same high frequency, leads to more early LTP and thereby triggers PRP synthesis. Also, the total actin-dependent volume shows a temporal summation of changes from multiple stimulations, which are then resulting in an increase of *V*_PSD_.

**Fig. 3.**
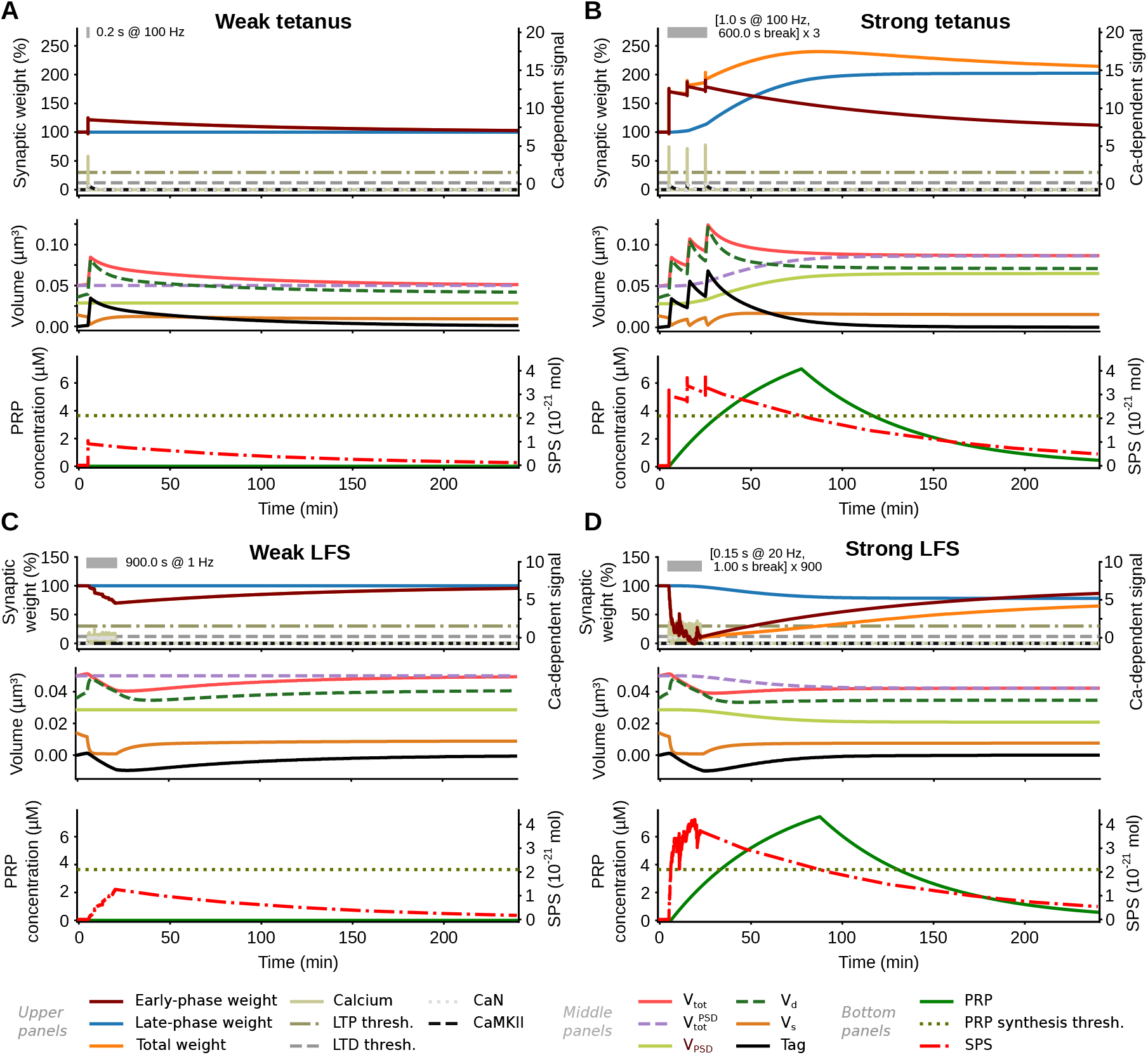
Temporal evolution of synaptic weights and relevant quantities in the full-integrated, calcium-based model under different stimulation protocols. Stimulation is conveyed by presynaptic spike activity. (A) Weak tetanization leads to e-LTP, (B) strong tetanization induces l-LTP, (C) weak low-frequency stimulation induces e-LTD, and (D) strong low-frequency stimulation yields l-LTD. *Upper panels:* Early-phase (brown, shifted for visualization purposes), late-phase (blue), and total weight (orange) of the individual synapse as well as calcium-dependent signals (calcium amount, CaN activity, and CaMKII activity, in normalized units). LTD and LTP thresholds are shown by dashed and dash-dot horizontal lines; CaMKII threshold *θ*_CaMKII_ = 5 not shown. Gray bar at the top: information on the time point, duration, and structure of the stimulation. *Middle panels*: Quantities of late-phase plasticity of the individual synapse (dynamic pool: dark green dashed, stable pool: ocher, total volume: pink, PSD volume: light green, 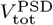: purple dashed; compare Fig. 2). *Bottom panels:* Quantities of neuron-wide processes – concentration of available PRP (dark green) and amount of the signal triggering PRP synthesis (SPS, red).

For low-frequency stimulation (Fig. 3C and D), we obtain calcium levels above the depression threshold but below the potentiation and CaMKII thresholds. Thus, actin nucleation is decreased and the overall actin-induced spine volume decays. For strong low-frequency stimulation, enough early-phase plasticity is expressed to trigger PRP synthesis, such that *V*_PSD_ decreases.

Thus, in summary, our results show that the difference between actin-dependent spine volume and PSD size also serves as synaptic tag in a biologically more detailed computational model under different experimentally inspired neuronal stimulation protocols, further supporting our hypothesis.

### Heterosynaptic plasticity via tagging-and-capture and the decay of the tag

As a next step, we investigated the behavior of our model under tagging-and-capture protocols for heterosynaptic plasticity. Specifically, we tested whether the proposed tag also shows the expected behavior if protein synthesis is triggered by stimulation at another synapse. For this, we apply weak and strong stimuli, with a time delay between them, at different synapses of the same neuron. The weak stimulus generates a synaptic tag at the stimulated synapse, but does not trigger PRP synthesis and thus no late-phase plasticity. Complementary to this, the strong stimulus at another synapse elicits PRP synthesis, such that newly synthesized PRPs can be used by the tag at the weakly stimulated synapse to form late-phase plasticity.

To test the according behavior of our fully-integrated model, we simulated a situation where the strong stimulus arrives before the weak stimulus. For this we find that the weak stimulus, which by itself could not cause changes of the late-phase weight (Fig. 3A), now indeed leads to a persistent increase in the PSD volume and, thus, the late-phase-weight (Fig. 4A, panels for synapse 2). Knowing that the full-integrated model captures this behavior, we systematically analyzed the determinants of heterosynaptic tagging-and-capture using our proof-of-concept actin-PSD model. We focused on the emergence of late-phase plasticity at the weakly stimulated synapse, considering that PRP synthesis is triggered by a strong stimulus at another synapse at a time before (strong-before-weak protocol, Fig. 4C) and after the weak plasticity event (weak-before-strong protocol, Fig. 4B, D). When the strong stimulus arrives after a weak tetanus (Fig. 4B), PRPs become available later than for the usual homosynaptic LTP (Figs. 2 and 3). At this time the total volume of the actin pools has almost decayed back to its initial value, and hence the adaptation of *V*_PSD_ needed to match this volume becomes smaller. Accordingly, we observe a decreased magnitude of late-phase LTP for a weak-before-strong tetanic protocol. Similarly, also for a weak LFS arriving before a strong stimulus (Fig. 4D), we observe a decreased magnitude of late-phase LTD. In contrast, for the strong-before weak protocol, PRP is available at the weakly stimulated synapse already at the time of stimulation such that the magnitude of late-phase changes is larger (Fig. 4C).

**Fig. 4.**
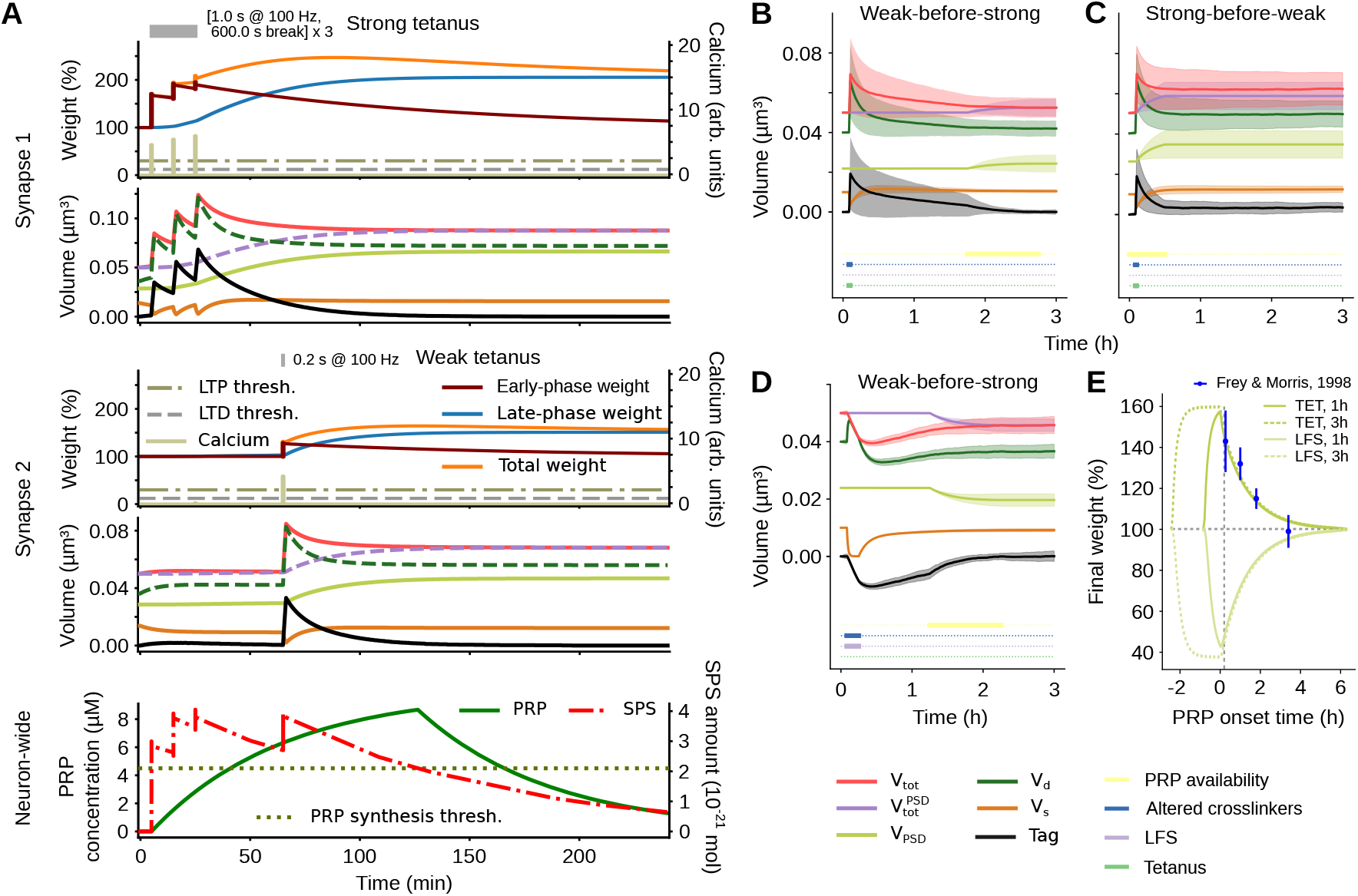
Heterosynaptic plasticity via tagging-and-capture. (A) Example of a strong-before-weak tetanization protocol in the full-integrated model (strong tetanus via first synapse starting at 300 s, weak tetanus via second synapse starting at 3900 s, cf. Fig. 3); (B) weak-before-strong tetanization in the late-phase only model; lines and shaded areas represent mean and standard deviation over 100 simulations, respectively; (C) same for strong-before-weak tetanization; (D) same for weak-before-strong low-frequency stimulation. (E) Final values of *V*_PSD_ in the late-phase-only model after weak tetanus (vivid color) and LFS (pale color), under systematic variation of the onset time of PRP availability due to a strong stimulus. Strongest plasticity occurs when PRP is available at the time of the weak stimulus. For PRP availability before the stimulus, the duration of PRP availability (1 h solid line, 3 h dashed line) determines the maximal time difference between strong and weak stimuli. The dashed vertical line marks the PRP availability used for the reduced model in the rest of the study. Experimental results [50] are shown for comparison (data points in blue).

We then systematically varied the onset time of PRP availability, finding that the maximal effect of both LTP and LTD is achieved if PRP synthesis starts together with the weak stimulus (Fig. 4E). If the strong stimulus occurs before the weak one, PRPs are available upon the weak stimulation and *V*_PSD_ quickly adapts to balance the mismatch with the actin volumes (Fig. 4C). However, PRPs need to be available until the stimulus-induced imbalance of the actin volumes and the PSD volume at the weakly stimulated synapses are fully equilibrated. Otherwise, the PSD dynamic will stop early and the rest of the equilibration is only achieved by dynamic pool changes (slower decay after *t* = 30 min in Fig. 4C). Hence, the duration of PRP availability determines how long before the weak stimulus the strong one can be started to yield maximal late-phase plasticity. If PRPs are available for one hour, maximal plasticity is achieved when the PRP synthesis starts immediately at the weak stimulus (Fig. 4E, solid lines). However, PRPs may be present for many hours [51], and therefore, we also tested a prolonged PRP availability of 3 hours (Fig. 4E, dashed lines). In this case, PRP synthesis could start up to 1.5 hours before the stimulus and still lead to maximal changes. On the other hand, if PRP synthesis starts after the stimulus (weak-before-strong), we observe the same gradual decay of the late-phase plasticity for both cases. Here, initially only the dynamic pool volume adapts to the PSD and decreases the imbalance. Consequently, less change of the PSD is needed to match the volume once PRPs are available, and the overall change of the PSD is smaller (Fig. 4B and E). By assessing the decline of this overall PSD change with the temporal distance between weak and strong stimulus, one can evaluate the lifetime of the tag (Fig. 4E). For our model, we selected the parameter values such that this relationship matches the one found in electrophysiological experiments (Fig. 4E blue data points based on [50]). For low-frequency stimulation and LTD, the resulting curves are qualitatively similar, but for decreased instead of increased weights (Fig. 4D and E).

Taken together, these findings show that the proposed synaptic tag can account for the capture of PRPs triggered by stimuli at other synapses while exhibiting a realistic decay over time.

### Impact of multiple subsequent stimuli at the same synapse

So far, we have only considered a single stimulus at each synapse. Yet, there may be multiple stimuli at the same synapse which jointly determine the synaptic tag and, thus, the long-term plasticity of the synapse. Hence, as a last step, we evaluate the response of our actin-PSD model to multiple plasticity-inducing stimuli. To reduce complexity, we restrain our analysis to the late-phase-only plasticity model.

First, we tested how consecutive tetanic and low-frequency stimuli would influence the weights in our model. We find that for a sufficiently small inter-stimulus interval (e.g., 5 minutes in Fig. 5A), there will be no late-phase plasticity, indicating that the tags from LTP and LTD events cancel each other out. However, for longer inter-stimulus intervals we observe that late-phase potentiation re-emerges (Fig. 5B), which is consistent with electrophysiological [52] and spine volume data [53] on tag resetting. Thus, we conclude that our model can also reproduce the late-phase plasticity produced by two subsequent, interacting stimuli as expected from tag resetting experiments.

**Fig. 5.**
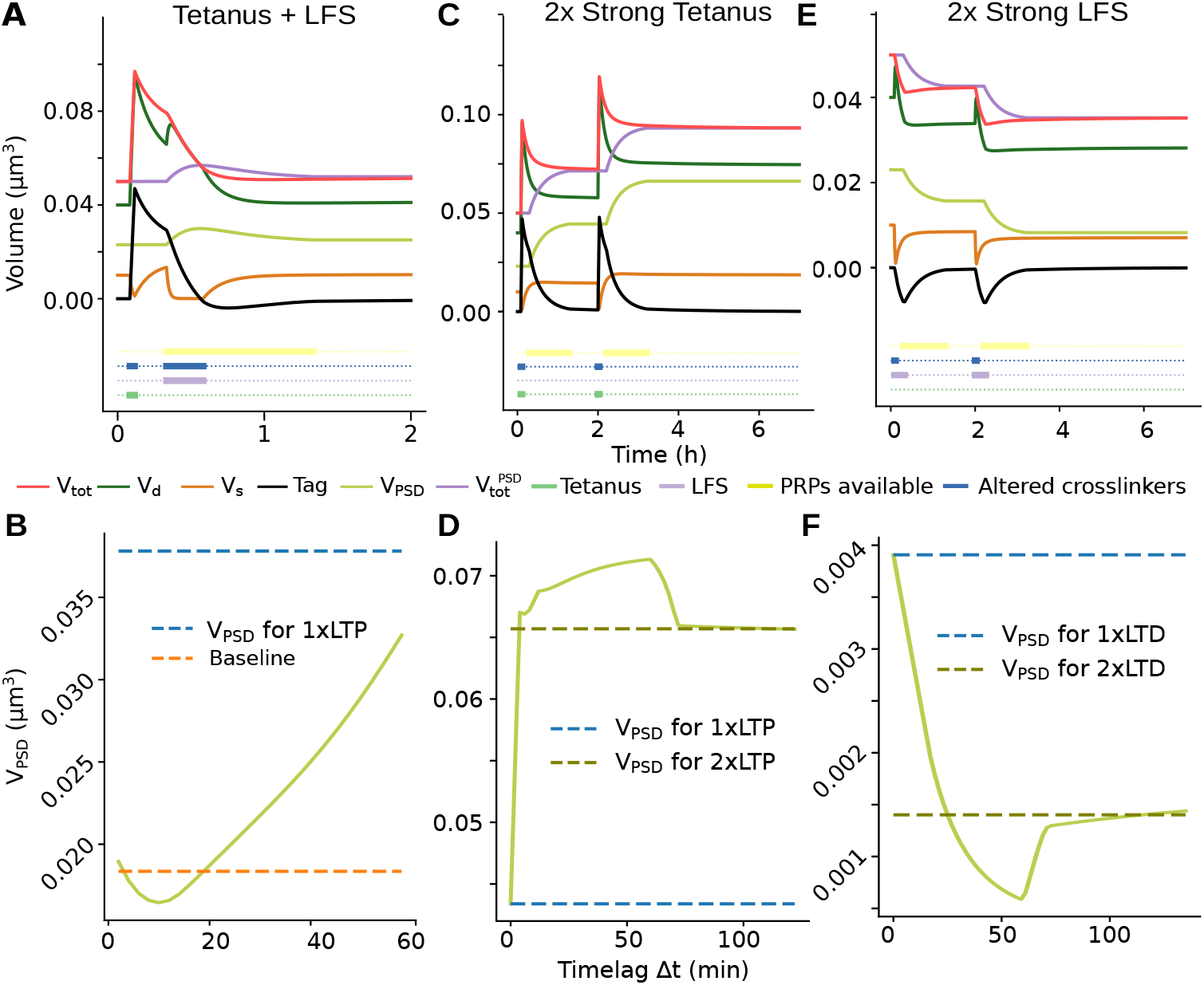
Impact of two temporally separated stimuli at one synapse. (A) Tag resetting: a strong LFS 5 min after a strong tetanus stimulus blocks the late-phase LTP effect. (B) Systematic analysis of the tag resetting effect depending on the LTP-LTD temporal lag, which reveals that tag resetting only works within a short time window after tag setting. (C) The effect of two strong tetanus stimuli accumulates linearly when they are sufficiently separated (here by 2 hours). (D) Dependence of the plasticity effect (i.e., of the final value of *V*_PSD_) on the temporal lag between two strong tetanus stimuli. (E) The effect of two strong LFS stimuli (again separated by 2 hours) accumulates linearly as well. (F) Dependence of the plasticity effect on the temporal lag between two strong LFS stimuli.

Finally, as a prediction of our model, we tested whether the lasting late-phase changes from multiple LTP- or LTD-inducing stimuli would sum linearly and how the timing between the stimulation events would influence this. For this, we first applied two high-frequency (tetanic) stimuli separated by a defined time interval (e.g., 2 h as shown in Fig. 5C) and evaluated the PSD size after all quantities had equilibrated. We compare these with the result of a single LTP event (blue dashed line in Fig. 5D) and of two consecutive but sufficiently separated LTP events (green dashed line, Fig. 5D). We observe that, for inter-stimulus intervals of a few minutes, the resulting volume and PSD size are smaller than for two independent events (sub-linear summation), indicating an occlusion of LTP (see [54] for experimental evidence of this). After that, however, we observe a super-linear summation of the LTP events for inter-stimulus intervals of up to around 75 minutes. After this interval, the PRP availability of the first stimulus has vanished and there is no influence on the second stimulus such that the results match those of two far separated LTP-events (linear stacking).

We further assessed how the temporal spacing between two low-frequency stimuli influences LTD (Fig. 5E and F). Similar as for LTP, we observe weaker LTD (sub-linear summation) for inter-stimulus-intervals up to around 45 minutes and stronger LTD (super-linear summation) in a time window from 45 to 75 minutes after the first stimulus (Fig. 5F).

Taken together, most importantly, our model predicts that around an hour after a first stimulus has been received, a repeated stimulus causes a stronger plasticity effect. This may relate to psychological effects such as spaced learning [55].

## Discussion

To provide a mechanistic description of the synaptic tag, we have derived a biophysically interpretable reaction rate model of actin and the PSD, which are the key players in structural and functional synaptic plasticity. Our model can reproduce experiments not only on homosynaptic late-phase LTD and LTP but also on heterosynaptic synaptic tagging-and-capture experiments. In summary, we conclude from our findings that the mismatch between the spine volume induced by actin dynamics and the volume expected from the PSD size constitutes a viable candidate implementation for the synaptic tag.

In our model, the tag is a continuous quantity that decays in the absence of PRPs due to the adaptation of the dynamic actin pool, enabling it to reproduce the graded decay of the plasticity outcomes in the weak-before-strong paradigm (Fig. 4E). Such a continuous decay would also be observable in other models that employ continuous tag variables [20, 21]. However, in these models the decay occurs exponentially over time, whereas the non-linearity introduced by the dynamic actin concentration term in our model would entail a non-exponential decay. Models that feature a continuous tag but no tag dynamics between plasticity events [56] can likely not reproduce the experiments. On the other hand, there are also models in which the tag is a discrete state [16, 17] and decays due to fixed transition rates. When considering a population of synapses [5] – as the weak-before-strong experiments do – also these models would exhibit an exponential decay of plasticity outcomes. Hence, to distinguish continuous and discrete tag models, further experiments on the tag decay at the single synapse level are needed.

Along this line, despite its biophysical motivation and matching with experimental data, our tagging mechanism remains an assumption of the model, which can, however, make experimentally testable predictions. The most straightforward approach would be to measure the correlation of PSD size and spine volume in basal state, stimulate with an LTP protocol and then again track these measures for individual spines (compare [31, 38]). The deviation from the basal correlation a few minutes after LTP – corresponding to the here-proposed synaptic tag – should then predict the change in PSD size and spine volume long after the LTP stimulus. Sufficient temporal resolution of the stimulation time-point would require the use of glutamate uncaging or electrical stimulation, and a better disentanglement of tag and early effects of consolidation could be achieved by blocking PRP synthesis until the first measurement. Still, such experiments would test a correlation between the PSD-size-and-spine-volume mismatch and PSD-dependent late-phase plasticity, but not a causation. To show the latter, one would have to artificially induce or remove a PSD-volume mismatch without changing the dynamics corresponding to other candidate mechanisms for tagging. Photoactivatable molecular constructs could be a promising approach to this. While such constructs have been successfully applied to study synaptic plasticity [57, 58], studies so far did not test constructs that target specifically actin dynamics (see, e.g. 59, 60), on which our long-lasting volume increases rely. However, one should be aware that actin interacts with a broad variety of partners and volume-change alone could alter the concentrations and thus the dynamics of all molecules within the spine. Thus, even if volume changes are a sufficient tagging condition, the tag might still be a complex structural state that comprise downstream effects of volume changes, so that testing a direct causation between PSD-spine-volume mismatch and PSD growth will remain challenging.

Another open question is the biophysical mechanism through which altered actin dynamics and spine size influence PSD growth. One mechanism that could account for this is that actin-driven increases in spine size let the membrane put less pressure onto the periphery of the PSD. This allows the PSD to utilize available PRPs to grow out along the periphery preserving its layered structure. We tested the former hypothesis in a more detailed biophysical simulation and found that this mechanism is indeed plausible [61]. Yet, apart from structural changes of the PSD, further processes such as persistently altered AMPA-receptor trafficking might play a role in consolidating synaptic changes [62].

Other predictions of our proof-of-concept model, which could be tested experimentally, concern the interaction of multiple, temporally adjacent stimuli (cf. [54, 55]). Most established phenomenological models of synaptic plasticity do not include interaction of consecutive stimuli on a timescale of minutes (cf. [63–65]). Yet, it can be very beneficial for neural systems: On the one hand, the adaptation of synaptic weights needs energy, and it is likely that the brain would employ plasticity mechanisms that are energy-efficient [66]. Recent research has shown that this energy-efficiency can be achieved by having a fast plasticity mechanism with small energy cost that integrates over a few stimuli and a slow plasticity mechanism with larger energy cost that adapts the synaptic weights only if needed [67]. In our model the fast process would be implemented by the structural changes that induce a continuous synaptic tag which is manifested by the imbalance between actin volumes and PSD in our model. Through multiple stimuli, this imbalance can be enlarged or shrinked and even pushed to the opposite sign. This process is likely energy-efficient as it is sufficiently implemented by slight tweaks of the local actin dynamics. The slow plasticity mechanism, given by the adaptation of the PSD, then transfers the tag into actual functional changes. This can be considered to be less energy-efficient as it involves *de novo* protein synthesis and/or transport of molecules (such as proteins or mRNA) from the soma. On the other hand, two-phase plasticity mechanisms have also been proposed to enhance storage capacity in neural networks [68–71]. In contrast to earlier plasticity models that often assume bistable late-phase weights, a line attractor, as considered here, may improve storage capacity by allowing for a continuous range of synaptic weights. For example, it can facilitate the storing of overlapping ensembles of neurons [72], where simpler bistable models would suffer from interference due to their inability to integrate multiple increases in synaptic weight (also see our investigations in Fig. 5). In theory [73], the line attractor can enable networks to store memory representations with an optimized set of weights, mitigating catastrophic forgetting and improving storage capacity.

Finally, the implementation of our full-integrated model with the Arbor simulator (see Methods) allows to seamlessly deploy the model in future network simulations with single- or multi-compartment neurons (cf. [74]). In particular, our own previous network models [20, 72, 75], which have considered a simpler model of synaptic tagging-and-capture combined with the same calcium-based model of early-phase plasticity that we used here, may thus easily be extended to account for the biophysically more realistic actin-based formulation that we have presented here. This may provide more detailed insights into the impact of multiple stacked learning stimuli as mentioned above (cf. [72]), the impact of the timing of PRP synthesis at the network level (possibly depending on neuromodulators, cf. [75]), as well as the priming of memory representations on timescales of the synaptic tag [72]. The Arbor simulator is explicitly built for multi-compartment neuron models and therefore allows for simulating detailed neuron morphologies and the respective propagation of molecules and currents [76]. Thus, also the effects of PRP transport from the nucleus or dendritic translation sites and their effect on local heterosynaptic plasticity could be studied using the provided implementation (cf. [56, 77]). Furthermore, with few adjustments, also other network models featuring synaptic tagging-and-capture ([18, 56, 70], see [78] for a review) may be improved by adding our proposed biophysical late-phase mechanisms.

## Methods

### Actin-based model of tagging-and-capture

Long-lasting synaptic alterations due to synaptic tagging-and-capture are known to depend on dynamic actin [41, 85]. Changes in actin dynamics affect spine geometry, which is followed by alterations of postsynaptic density [38]. These, in turn, correlate with the number of bound AMPA receptors and, thus, the synaptic weight. Following this reasoning, we model tagging-and-capture based on the amount of dynamic, fast-treadmilling actin (via the volume *V*_d_ induced by the dynamic pool), the amount of cross-linked, stable actin (via the volume *V*_s_ induced by the stable pool), and the size of the postsynaptic density (via the volume *V*_PSD_ induced by the postsynaptic density).

The dynamics of the system are given by:

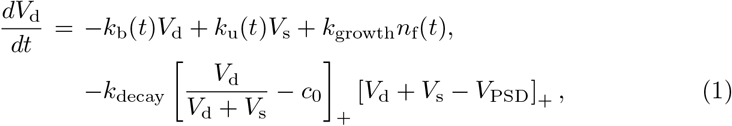

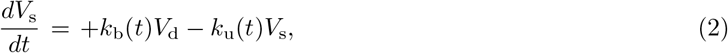

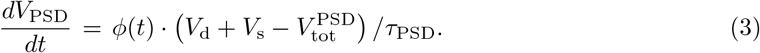

Here, *k*_b_ and *k*_u_ are the crosslinker binding and unbinding rates, *k*_growth_ and *k*_decay_ the rates for dynamic pool growth and shrinkage, *ϕ*(*t*) the PRP availability, *τ*_PSD_ the PSD reorganization timescale, [*x*]_+_ = (*x* + |*x*| )*/*2 the rectification function, and *n*_f_ the number of active polymerization foci. More detailed simulations of the actin network suggest that *n*_f_ fluctuates heavily because new foci continuously emerge while old foci die out [23, 26]. To approximate this, we here determine *n*_f_ by a birth-death process, with a birth rate *γ* and a death rate *µ*. The same simulations also showed that the number of foci is a good predictor for volume change [23], such that we model volume growth proportional to *n*_f_ although the relationship between spine geometry and actin dynamics is much more complicated (c.f. [23, 26, 33])

While

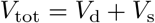

represents the total volume of the spine determined by the actin dynamics,

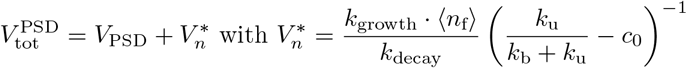

is the steady-state volume that can be expected for the current PSD size with dynamic actin in its basal state (which induced the correction 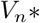; see Appendix). Here, ⟨.⟩ denotes the temporal average (in basal state). Note, in the absence of dynamic actin foci (*n*_f_ = 0), 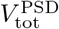 equals *V*_PSD_.

### Time-dependent parameters during LTP and LTD induction

During plasticity, the crosslinker binding and unbinding as well as the birth and death rate of actin foci are altered as described in the following. All constant parameters are reported in Table 1.

**Table 1.** Parameter values and sources for both the late-phase model and the other parts of the full-integrated model.

| Param. | Value | Description | Reference |
| --- | --- | --- | --- |
| $k_{\text{growth}}$ | $135 \cdot 10^{-6} \text{ s}^{-1} \mu\text{m}^3$ | growth rate of the dynamic actin pool | |
| $k_{\text{decay}}$ | $1/380 \text{ s}^{-1}$ | decay rate of the dynamic actin pool | [5] |
| $\tau_{\text{PSD}}$ | $380/0.255 \text{ s}$ | adaptation rate of the PSD (with PRPs) | [38] |
| $c_0$ | 0.725 | basal fraction of dynamic actin | |
| $\tau_c$ | 0.0488 s | calcium timescale | [19, 20, 49] |
| $c_{\text{pre}}$ | 1 | calcium increase per presynaptic spike | [19, 20, 49] |
| $c_{\text{post}}$ | 0.2758 | calcium increase per presynaptic spike | [19, 20, 49] |
| $t_{\text{cad}}$ | 0.0188 s | delay of calcium increase with respect to presynaptic spike | [19, 20, 49] |
| $\theta_p$ | 3.0 | potentiation threshold | [19, 20] |
| $\theta_d$ | 1.2 | depression threshold | [19, 20] |
| $\theta_{\text{CaMKII}}$ | 5.0 | threshold for CaMKII activity | |
| $\gamma_p$ | 1645.6 | early-phase potentiation rate | [49] |
| $\gamma_d$ | 313.1 | early-phase depression rate | [49] |
| $\tau_h$ | 688.4 s | early-phase weight timescale | [49] |
| $\sigma_{\text{wn}}$ | 2.90436 mV | scaling factor for fluctuations of early-phase plasticity | [49, 72] |
| $\tau_{\text{CaMKII}}$ | 120 s | timescale of CaMKII deactivation | [79, 80] |
| $\tau_{\text{CaN}}$ | 120 s | timescale of CaN deactivation | [80] |
| $\theta_\phi$ | 2.10037 mV | PRP synthesis threshold | [19] |
| $\tau_\phi$ | 3600 s | PRP timescale | [16, 19] |
| $\phi_{\text{max}}$ | 10.0 $\mu\text{mol/l}$ | equilibrium (maximum) concentration of PRPs under ongoing synthesis | [16, 19, 74] |
| $f_{\text{int}}$ | 0.11/$\mu\text{mol}$ | scaling of the integration of PRPs into the PSD | [74] |
| $c_w$ | 4.20075 mV | constant determining the initial value and dynamics of the total synaptic weight | [49, 72] |
| $c_{\text{late}}$ | 0.8 | late-phase fraction of the total weight | [19, 20] |
| $\tau_{\text{mem}}$ | 10 ms | membrane time constant | [63, 81] |
| $\tau_{\text{syn}}$ | 5 ms | synaptic time constant | [63, 81, 82] |
| $t_{\text{axd}}$ | 3 ms | axonal transmission delay | [81, 83] |
| $t_{\text{ref}}$ | 2 ms | absolute refractory time | [63, 84] |
| $V^{\text{rev}}$ | -65 mV | reversal (equilibrium) voltage | [63] |
| $V^{\text{th}}$ | -55 mV | threshold voltage for firing | [63] |
| $V^{\text{reset}}$ | -70 mV | reset voltage | [63] |

**Table 2.** Time-dependent changes of the crosslinker and actin foci dynamics after stimulation. The reported ⟨*n*_f_⟩ is used in simulations, where the birth-death-process is replaced by a constant *n*_f_-value and tuned to yield results comparable to the stochastic simulations.

| Stimulus | Birth $\gamma$ | Death $\mu$ | $\langle n_f \rangle$ | $k_b$ | $k_u$ | Interval | PRPs available |
| --- | --- | --- | --- | --- | --- | --- | --- |
| Basal | 0.02 s<sup>-1</sup> | 0.5 s<sup>-1</sup> | 0.04 | 1/2480 s<sup>-1</sup> | 1/620 s<sup>-1</sup> | - | - |
| Weak tetanus | 0.45 s<sup>-1</sup> | 0.5 s<sup>-1</sup> | 3.12 | 1/55000 s<sup>-1</sup> | 1/55 s<sup>-1</sup> | [0 s, 60 s] | - |
| Strong tetanus | 0.45 s<sup>-1</sup> | 0.5 s<sup>-1</sup> | 3.12 | 1/55000 s<sup>-1</sup> | 1/55 s<sup>-1</sup> | [0 s, 120 s] | [900 s, 4300 s] |
| Weak LFS | 0.0 s<sup>-1</sup> | 0.05 s<sup>-1</sup> | 0.0 | 1/55000 s<sup>-1</sup> | 1/55 s<sup>-1</sup> | [0 s, 600 s] | - |
| Strong LFS | 0.0 s<sup>-1</sup> | 0.05 s<sup>-1</sup> | 0.0 | 1/55000 s<sup>-1</sup> | 1/55 s<sup>-1</sup> | [0 s, 900 s] | [900 s, 4300 s] |

After stimulation, the actin cytoskeleton can undergo highly dynamic changes as actin-binding proteins change their concentrations and binding rates. For example, cofilin gets activated after an LTP-inducing stimulus, which leads to a massive severing and, in turn, produces an abundance of free actin and creates new foci [28, 31]. Moreover, crosslinkers unbind from the filaments [27, 28, 31]. Although not so well documented, the latter may also occur for LTD [41, 45, 46].

In order to study our actin-PSD model (Eqs. 1–3) during plasticity, we first implemented time-dependent parameters in the form of step functions, which switch their value at given time-points after a putative LTP or LTD event. We used the following alterations, which are summarized in Table 2:

#### Weak and strong stimulations

In general, strong stimuli induce late-phase plasticity, while weak stimuli only induce early-phase effects. Thus, for weak stimuli, the duration of the rate alterations is smaller and *ϕ*(*t*) is kept at zero throughout the experiment. In contrast, for strong stimuli, we set the PRP availability to *ϕ* ≡ 1 for 60 minutes starting 15 minutes after the stimulus (if not stated otherwise; also cf. PSD changes in [38]).

#### Tetanic stimulation

Tetanic, i.e., high-frequency stimulation can induce LTP. Accordingly, immediately after a tetanic stimulation event, the birth rate of the stochastic process is significantly increased for the typical duration of a tetanization protocol (60 s/120 s for weak and strong, respectively). This corresponds to an increase of cofilin in the spine, which has experimentally been observed in the early phase of LTP [31]. Also, within the first minutes of LTP, crosslinkers unbind from actin and/or leave the spine [31, 45, 86]. Therefore, in the model, the crosslinker binding and unbinding rates *k*_b_ and *k*_u_ rates are increased to allow faster exchange between the two pools, shifting their balance towards unbinding, which results in a smaller stable actin pool.

#### Low-frequency stimulation (LFS)

LFS can give rise to LTD. Accordingly, in our model, we set the birth rate of actin foci to zero and adapt the death rate for the duration of the stimulus (600 s/900 s). The crosslinker binding rates are subject to the same transformation as in LTP, but in this case, the duration of the effect is prolonged to remain for the duration of the LTD stimulus.

### Parameter estimation based on experimental data

To find parameters that match experimental characterizations of synapses, we used three different datasets: (1) volume time courses after glutamate uncaging inducing LTP [42] and (2) LTD [48], as well as (3) electrophysiological data from the weak-before-strong protocol [50]. For each datapoint, we calculated the squared error of the relative volume changes, normalized by the size of the experimentally observed volume change. We weighted the errors of the 3 datasets with *w* = (1, 2, 10)^*T*^ to account for the imbalance between the number of data points, and calculated the sum over all these weighted normalized errors. We then minimized this error. For numerical stability, we used the inverse of the rates as well as the product of covariant parameters as free parameters (with their bounds in brackets):

- the timescale for crosslinker unbinding 1*/k*_u_ ∈ [300 s, 1500 s]
- the dynamic pool decay timescale 1*/k*_decay_ = 240 s (varied in step 2 only, see below)
- the ratio between dynamic pool decay and PSD adaptation timescales ∈ [0.05, 0.5]
- the product (mean) number of active foci and the growth rate 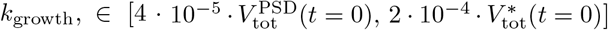
- same product during 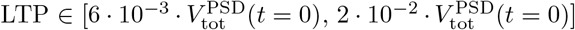
- the offset fraction of the dynamic pool *c*_0_ ∈ [0.7, 0.75]

For minimization, we used a two-step procedure:

1. *Bayesian optimization:* Using the Bayesian Optimization package [87], we scanned for local optima in the above specified parameter regime by using 200 initial samples and 50 samples from the Optimizer.
2. *Minimizer* The resulting best parameter set was then passed to a gradient descent optimizer (Nelder-Mead from scipy optimize) in order to reach the local minimum.

Here, the parameter changes were bounded within 10% of the initial value. Throughout the process, rates were rounded to multiples of 5 s.

After that, the ratio between binding and unbinding timescales in basal conditions was set to 1/4 to obtain a 20% stable pool fraction [29], while under stimulation, the ratio was set to 1/1000 to effectively dissolve the stable pool. The collinearity of *n*_f_ and *k*_growth_ was resolved by fixing the basal *n*_f_ to 0.04 corresponding to a focus nucleation rate of *γ* = 1*/*50 s^−1^ and death rate of *µ* = 1*/*2 s^−1^.

The robustness of the simulation results with respect to changes in the fitted parameters is reported in Appendix B.

### Fully integrated model with calcium-based early-phase plasticity and PRP synthesis

In order to study its interaction within neurons and neural circuits, we coupled our model of late-phase plasticity to established models for neuronal activity, calcium-based early-phase plasticity, and PRP synthesis (cf. [19, 20, 78]). Large parts of the formulation below thus follow previous descriptions [20, 72]. Note that as compared to [20, 72], we use here a transformed formulation of the early- and late-phase dynamics where the baseline of the early-phase weights is *h*(*t* = 0) = 0 and the baseline of the late-phase weights is *z*(*t* = 0) = *c*_*w*_. This formulation is equivalent to the one in the previous studies, but may be easier to interpret from a biological perspective (the baseline of the total weight is here set by the stable late-phase weight).

### Calcium

LTP and LTD are known to rely on calcium signaling both on a morphological and functional level [42, 43, 49]. Therefore, in our model, the calcium level *c*_*ji*_(*t*) at the postsynaptic site of a synapse from neuron *i* to neuron *j* is the major factor to induce plasticity. Following [49], the calcium level is increased by *c*_pre_ for every presynaptic spike at time *t*^*n*^ and by *c*_post_ for every postsynaptic spike at time *t*^*m*^, and it evolves according to:

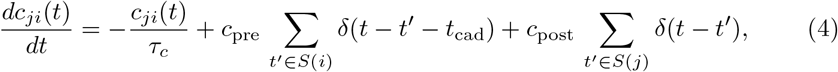

where *S*(*i*) and *S*(*j*) mark the set of spike times of neurons *i* and *j*, respectively, δ(·) is the Dirac delta distribution, *τ*_*c*_ is a time constant, and *t*_cad_ accounts for the delayed increase of calcium for presynaptic spikes.

### Early-phase plasticity

Early phase-plasticity is based on altered receptor trafficking driven by the calcium concentration in the postsynaptic spine [36, 88]. We model this by an early-phase weight *h*(*t*) following a calcium-based plasticity rule with two thresholds for depression and potentiation, respectively [19, 20, 49]. The dynamics are described by:

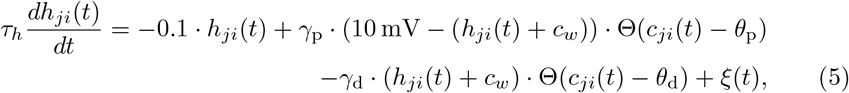

where Θ(·) is the Heaviside step function, *c*_*w*_ is a constant that determines the initial value and dynamics of the total synaptic weight, *τ*_*h*_ is the calcium decay time constant, *c*_*ji*_(*t*) is the calcium concentration at the postsynaptic site, and *ξ*(*t*) is a noise term (see below).

The first term on the right-hand side describes the relaxation of the early-phase weight to zero, the second term describes the induction of early-phase LTP with rate *γ*_p_ (requiring that the calcium amount is above the threshold *θ*_p_), and the third term describes the induction of early-phase LTD with rate *γ*_d_ (requiring that the calcium amount is above the threshold *θ*_d_). As the latter two terms are multiplied by (*h*_*ji*_(*t*) + *c*_*w*_) and (10 mV − (*h*_*ji*_(*t*) + *c*_*w*_)), the early weight is soft-bounded to an interval [− *c*_*w*_, 10 mV − *c*_*w*_]. Finally, the noise term depends on Gaussian white noise Γ(*t*) with mean zero and variance 1*/dt*, and is given by (cf. [49])

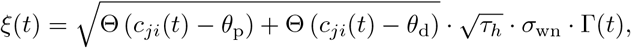

with the factor *σ*_wn_ scaling its standard deviation.

### Synthesis of PRPs

Synthesis of plasticity-related products (PRPs), in particular, of proteins, has been found to occur in the soma as well as locally in dendrites. In our full-integrated model, we describe PRP dynamics as a process of synthesis and decay that may occur neuron-wide or in a locally clustered manner. To determine the onset of PRP synthesis, we define a “signal triggering PRP synthesis” (SPS) as the absolute deviation of the early-phase weights from their stationary value 0 summed over all considered incoming synapses (cf. [19]):

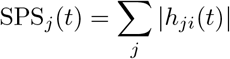

If this is larger than a threshold *θ*_*ϕ*_, then the amount of PRP *ϕ*_*j*_(*t*) increases to a maximum value *ϕ*_max_, whereas it decays to the baseline zero otherwise:

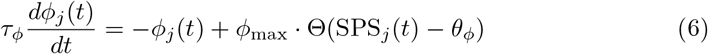

Here, the timescale of the dynamics is given by the constant *τ*_*ϕ*_.

### Coupling of early-phase dynamics to the actin-PSD model of late-phase plasticity

The changes in the dynamics of actin-binding proteins during LTP and LTD are induced by calcium-dependent signaling cascades [28]. In particular, the signaling chain depending on calcineurin (CaN) is activated by intermediate calcium levels during LTD, whereas the signaling chain downstream of CaMKII is typically activated by very high calcium levels and triggers LTP in further downstream signaling. Following this, in our integrated model, CaMKII and CaN are set to CaMKII ≡ 1.0 and CaN ≡ 1.0 whenever the calcium concentration fulfills *c*(*t*) *> θ*_CaMKII_ and/or *c*(*t*) *> θ*_d_, respectively (cf. Eq. 4).

Otherwise, the concentrations of CaMKII and CaN decay exponentially following

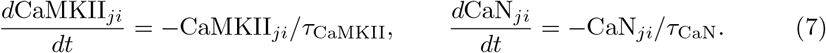

Altered dynamics for the crosslinkers and the birth-death process are applied if one or both of the signals are above a threshold of 0.5. For simplicity, we do not explicitly simulate the birth-death process for actin foci in the full-integrated model, but instead use the expected mean values for *n*_f_ as given in Table 2. Thus, *k*_b_, *k*_u_ and *n*_f_ for each synapse are set to the values for potentiation from Table 2 when CaMKII-signaling is on (CaMKII_*ji*_ *>* 0.5), to the values for depression when only CaN-signaling is on (CaMKII_*ji*_ *<* 0.5 and CaN_*ji*_ *>* 0.5), and to the basal values otherwise.

Finally, in our integrated model, we use the PSD-dependent volume *V*_PSD_ as a proxy for the late-phase weight *z* (cf. [20, 72]) in the following way:

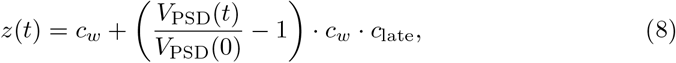

where *c*_*w*_ is a constant determining the unscaled initial weight and the weight dynamics, and *c*_late_ determines the ratio by which the late-phase weight enters the total weight. Note that we here use a linear relationship between *V*_PSD_ and the synaptic weight, whereas the real relationship can be assumed to be highly nonlinear as *V*_PSD_ is not the size or volume of the PSD itself, which correlates with the synaptic currents.

Also the relation between PSD size and weight is likely nonlinear due to cooperative effects [36, 89]. Yet, the relations likely remain monotonic, such that the results from the assumed linear relationship will hold at least qualitatively.

The late-phase weight is then added to the early-phase weight *h* to yield the total weight of the synapse:

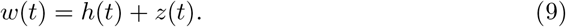

Note that as mentioned above, this is a transformed formulation as compared to [20, 72].

### Neuronal dynamics and connectivity

To model the neuronal membrane dynamics, we employed the leaky integrate-and-fire formalism, given by

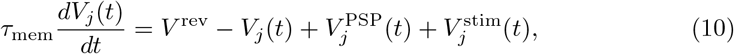

for the neuron with index *j*. Here, *V* ^rev^ is the reversal potential, *τ*_mem_ is the membrane time constant, 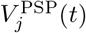 is the sum of postsynaptic potentials, and 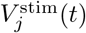 accounts for the external stimulation (see the following subsection). With the synaptic weights *w*_*ji*_ (see Eq. 9) and the set *S*(*i*) of the presynaptic spike times, the postsynaptic potential is given by:

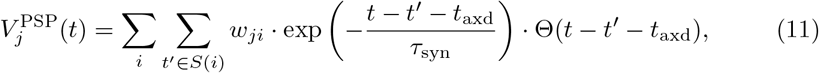

where *t*_axd_ is the axonal delay time, *τ*_syn_ is the synaptic time constant, and Θ( ·) is the Heaviside step function. A spike is triggered if the neuronal membrane voltage *V*_*j*_(*t*) crosses the threshold *V* ^th^, following which the membrane voltage is set to *V* ^reset^, where it remains for the refractory time *t*_ref_ (see Table 1 for the parameter values).

For the single-synapse paradigms shown in Fig. 3, we used two such neurons, connected by one synapse described by our integrated model of early- and late-phase synaptic plasticity with the calcium traces driven by the pre- and postsynaptic neuronal spikes. The presynaptic neuron was subject to the stimulation protocols described in the following subsection.

For the strong-before-weak paradigm in Fig. 4, we used three neurons {0, 1, 2} with the connections 1 → 0 and 2 → 0. In this context, neuron 1 received strong stimulation and neuron 2 received weak stimulation, as detailed in the next subsection.

Note that our freely provided Arbor program code can be used to explore the dynamics of our plasticity model in a wide variety of paradigms, including large-scale neural circuits.

### Stimulation protocols

To induce l-LTP, l-LTD, e-LTP only, and e-LTD only, we chose classical well-established stimulation protocols (cf. [19, 20, 44]). As described in the previous subsection, we conveyed these protocols by forcing a presynaptic neuron to spike with Poisson-distributed spike times, which can be used to model the firing of a neuron at a certain average rate (cf. [63]). The details of the protocols are described in the following.

#### Strong tetanic stimulation (STET)

This protocol serves to induce l-LTP. It consists of three pulses of 100 Hz stimulation, each lasting for 1.00 s. The pulses are separated by breaks of 600 s.

#### Weak tetanic stimulation (WTET)

This protocol serves to induce e-LTP. It consists of one pulse of 100 Hz stimulation, lasting for 0.20 s.

#### Strong low-frequency stimulation (SLFS)

This protocol serves to induce l-LTD. It consists of 900 pulses of 20 Hz stimulation, each lasting for 0.15 s. The pulses are separated by breaks of 1.00 s.

#### Weak low-frequency stimulation (WLFS)

This protocol serves to induce e-LTD. It consists of one long period of 1 Hz stimulation, lasting for 900 s.

### Software

For the late-phase-only model, we used a custom Python implementation that was designed to enable easy switching between simulation paradigms.

The full-integrated model was implemented using the Arbor simulator library [74, 90], which is particularly well suited for the efficient execution of network simulations with multi-compartment neurons on high-performance computing hardware, and will thus allow for future network simulations employing our model.

We will make our simulation code freely available upon publication of this article.

## Supplementary information

**Appendix A:** Fixed-point and stability analysis for the late-phase model

**Appendix B:** Parameter sensitivity

## Declarations

### Funding

This work was supported by the German Research Foundation (Deutsche Forschungsgemeinschaft, DFG) through grants SFB1286 (projects C01, C03 and Z01).

### Conflict of interest

There is no conflict of interest

### Data and Code availability

Data and Code for the reproduction of these results are available under [91].

### Author contribution

Conceptualization: FN, JL, CT, MF; Formal Analysis: FN, JL, MF; Funding acquisition: MF, CT; Investigation: FN, JL, MF; Methodology: FN, JL, CT, MF; Project administration: CT, MF; Software: FN, JL; Supervision: CT, MF; Visualization: FN, JL, MF; Writing – original draft: JL, MF; Writing – review & editing: FN, JL, CT, MF.

## Appendix A Fixed-point and stability analysis for the late-phase model

Here, we want to calculate the fixed points of our model under constant parameter values and analyze their stability. For this we first replace the stochastic birth-death-process that determines the number of foci with the current expected value, corresponding to the temporal mean ⟨*n*_f_⟩, to arrive at a non-stochastic dynamic system. We further simplify the calculation by defining *V*_tot_ = *V*_d_ + *V*_s_ and only investigating the dynamic of a two-dimensional system. When the crosslinker binding and unbinding are in equilibrium, the dynamic and stable pool can be obtained as fractions of *V*_tot_:

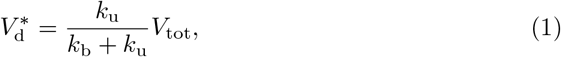

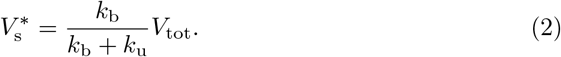

We arrive at

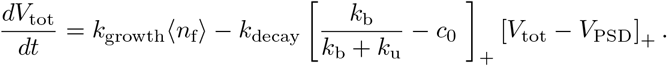

Assuming that both rectifications are in their linear regime, we obtain a fixed-point for *V*_tot_ that can be expressed as function of *V*_PSD_:

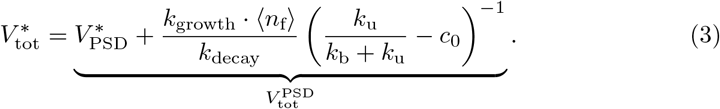

This constitutes a continuous line of fixed points in the *V*_tot_-*V*_PSD_-space. In order to compute the stability of these fixed points, we first compute the Jacobian matrix of our equations evaluated at the fixed point:

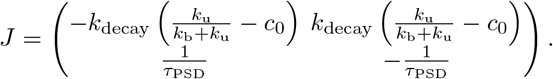

The resulting eigenvalues are *λ*_1_ = 0 and 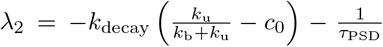. The eigenvector for *λ*_1_ is proportional to (1, 1)^*T*^ and thus aligns with the line of fixed-points. This proves that the fixed-point manifold defined by equation 3 forms a stable line attractor.

## Appendix B Parameter sensitivity

**Table 1.**
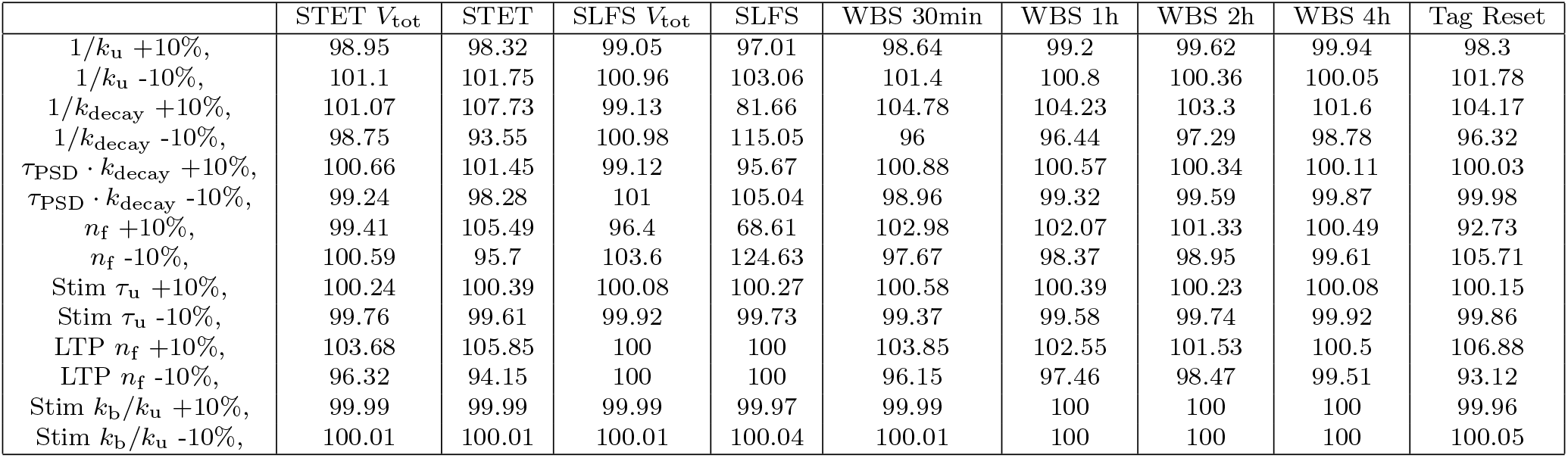
The final values of *V*_PSD_ and *V* tot (see column head) if the individual model parameters indicated in the rows are varied by 10% relative to their values with the model parameters used throughout the manuscript (in percent). For better comparability all parameters are either timescales (1 over rate) or unitless, as they were used for parameter fitting.

| | STET $V_{\text{tot}}$ | STET | SLFS $V_{\text{tot}}$ | SLFS | WBS 30min | WBS 1h | WBS 2h | WBS 4h | Tag Reset |
| --- | --- | --- | --- | --- | --- | --- | --- | --- | --- |
| $1/k_u +10\%$, | 98.95 | 98.32 | 99.05 | 97.01 | 98.64 | 99.2 | 99.62 | 99.94 | 98.3 |
| $1/k_u -10\%$, | 101.1 | 101.75 | 100.96 | 103.06 | 101.4 | 100.8 | 100.36 | 100.05 | 101.78 |
| $1/k_{\text{decay}} +10\%$, | 101.07 | 107.73 | 99.13 | 81.66 | 104.78 | 104.23 | 103.3 | 101.6 | 104.17 |
| $1/k_{\text{decay}} -10\%$, | 98.75 | 93.55 | 100.98 | 115.05 | 96 | 96.44 | 97.29 | 98.78 | 96.32 |
| $\tau_{\text{PSD}} \cdot k_{\text{decay}} +10\%$, | 100.66 | 101.45 | 99.12 | 95.67 | 100.88 | 100.57 | 100.34 | 100.11 | 100.03 |
| $\tau_{\text{PSD}} \cdot k_{\text{decay}} -10\%$, | 99.24 | 98.28 | 101 | 105.04 | 98.96 | 99.32 | 99.59 | 99.87 | 99.98 |
| $n_f +10\%$, | 99.41 | 105.49 | 96.4 | 68.61 | 102.98 | 102.07 | 101.33 | 100.49 | 92.73 |
| $n_f -10\%$, | 100.59 | 95.7 | 103.6 | 124.63 | 97.67 | 98.37 | 98.95 | 99.61 | 105.71 |
| Stim $\tau_u +10\%$, | 100.24 | 100.39 | 100.08 | 100.27 | 100.58 | 100.39 | 100.23 | 100.08 | 100.15 |
| Stim $\tau_u -10\%$, | 99.76 | 99.61 | 99.92 | 99.73 | 99.37 | 99.58 | 99.74 | 99.92 | 99.86 |
| LTP $n_f +10\%$, | 103.68 | 105.85 | 100 | 100 | 103.85 | 102.55 | 101.53 | 100.5 | 106.88 |
| LTP $n_f -10\%$, | 96.32 | 94.15 | 100 | 100 | 96.15 | 97.46 | 98.47 | 99.51 | 93.12 |
| Stim $k_b/k_u +10\%$, | 99.99 | 99.99 | 99.99 | 99.97 | 99.99 | 100 | 100 | 100 | 99.96 |
| Stim $k_b/k_u -10\%$, | 100.01 | 100.01 | 100.01 | 100.04 | 100.01 | 100 | 100 | 100 | 100.05 |

We here report the sensitivity of our results with respect to the fitted parameters. We varied each of these individually by 10% and tracked the outcome for *V*_tot_ and *V*_PSD_ 4 hours after the stimulation. We then report the relative size as compared to the original model parameters in Table 1.

The strongest influence is observed for changing the basal number of actin foci *n*_f_. For larger *n*_f_, we observe less LTP and more LTD, as the differences to the stimulation related number of foci decreases and increases, respectively. The value of the LTP induced number of foci is equally influential, but only for LTP. The relative influence of the timescales and their ratios is rather small.

We therefore conclude that our model is robust to changes of its parameters, while remaining sensitive to changes in the actin dynamics, which are providing the stimulus.

## Notes

### Competing Interest Statement

The authors have declared no competing interest.

### Summary of Updates

Updated discussion of alternative models and experimental predictions. Added robustness analysis.

